# Metabolomic characterization of *Epimedium sagittatum* bee pollen reveals a distinctive chemical profile

**DOI:** 10.1101/2025.09.29.679240

**Authors:** De-fang Niu, Hai-xin Chen, Cao-yang Lu, Cui-ping Zhang, Qiu-lan Zheng, Xiao-ling Su, Xiao-ming Fan, Yi-bo Luo, San-yao Li, Bin Yuan, Ping Liu, Fu-liang Hu

**Author notes:** Correspondence to*: Ping Liu, Jiangsu Agri-Animal Husbandry Vocational College, Taizhou 225300, China; E-mail address;Bin Yuan, College of Animal Sciences, Zhejiang University, Hangzhou 310058, China; E-mail address; Fu-liang Hu, College of Animal Sciences, Zhejiang University, Hangzhou 310058, China.

## Abstract

*Epimedium sagittatum* bee pollen (EBP) is a bee pollen product of a medicinal plant, but its chemical characteristics remain unclear. In this study, EBP was systematically characterized by biochemical analysis and UPLC–MS/MS, using *Brassica rapa* bee pollen (BBP), *Camellia sinensis* bee pollen (CBP), and *Epimedium* leaves for compari-son. A total of 1,073 secondary metabolites were identified in EBP, mainly flavonoids (330, 30.8%) and phenolic acids (165, 15.3%). EBP showed the highest total flavonoid content among the three bee pollen types (4.75 mg/g) and was clearly separated from BBP and CBP in multivariate analysis. EBP contained 19 unique metabolites, fewer than CBP (271), but these included several high-content flavonoid compounds, such as cacticin, brassicin, and tricetin-4′-methyl ether-3′-β-D-glucoside. Differential metabolite analysis identified 449 and 1,085 metabolites that differed between EBP and BBP, re-spectively, with flavonoid compounds forming the main differential class. Among the shared differential metabolites, 45 flavonoids were consistently higher in EBP, in-cluding kaempferol, tamarixetin, and sinensetin. Network pharmacology screening further suggested that flavonoid metabolites, especially kaempferol, tamarixetin, and sinensetin, deserve particular attention in future studies of EBP. Compared with Epimedium leaves, characteristic leaf flavonoids such as icariin and epimedins A–C were present at very low levels or were not detected in EBP. Overall, EBP is charac-terized by a distinct, relatively concentrated flavonoid chemical profile, distinct from both common commercial bee pollens and *Epimedium* medicinal tissues.

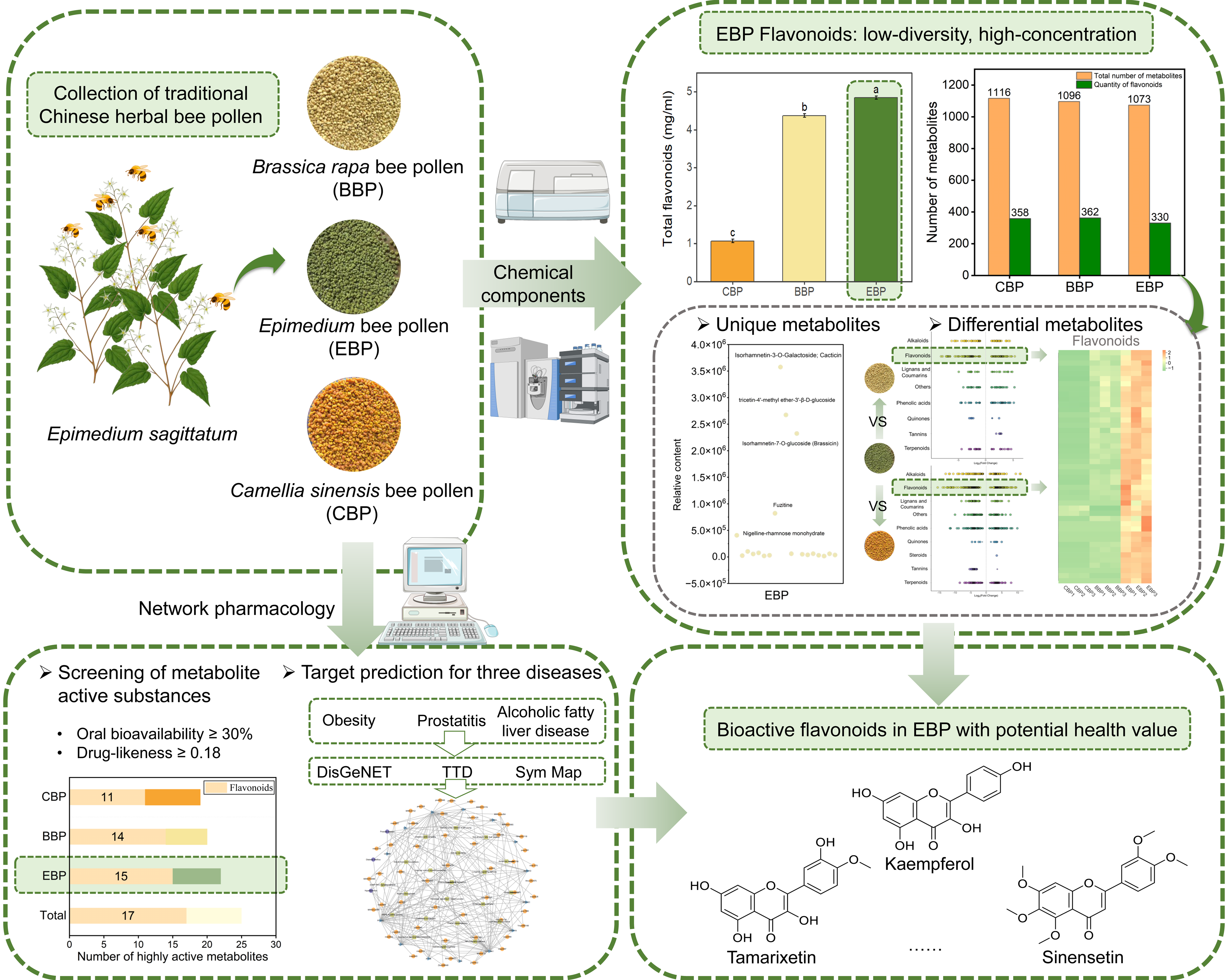

## 1. Introduction

Global consumption of dietary supplements and functional foods is rapidly increasing (<u>Di Martino 2025</u>). Bee pollen (BP) is an ideal candidate for such applications due to its rich content of bioactive phytochemicals, particularly flavonoids (<u>Giampieri, et al. 2022</u>). Widely commercialized BPs, such as *Brassica rapa* bee pollen (BBP) and *Camellia sinensis* bee pollen (CBP), exhibit well-documented anti-inflammatory and antioxidant properties (<u>Han, et al. 2023</u>; <u>Li, et al. 2023</u>). Recent studies consistently emphasize that botanical origin is a primary determinant of the biochemical complexity and nutritional profile of BP (<u>Bridi, et al. 2022</u>). Consequently, there is growing interest in BP derived from traditional Chinese medicinal herbs, such as *Codonopsis pilosula*, *Fagopyrum esculentum*, and *Schisandra chinensis*, which may offer enhanced therapeutic potential due to their unique secondary metabolites(<u>Huang, et al. 2017</u>; <u>Li, et al. 2024b</u>; <u>Niu, et al. 2021</u>).

However, evaluating the chemical characteristics of medicinal plant-derived BP requires careful consideration of tissue-specific metabolism (<u>Yan, et al. 2024</u>). Traditional medicines typically utilize vegetative tissues (e.g., leaves, stems, roots) (<u>Giménez-Bastida and Zieliński 2015</u>; <u>Luan, et al. 2021</u>). In contrast, pollen is a highly specialized reproductive tissue (<u>Rivest and Forrest 2019</u>). Because plant secondary metabolism exhibits strong tissue specificity driven by different physiological functions, the bioactive profile of the BP of a medicinal plant may not necessarily mirror that of its traditional medicinal parts (<u>Iiyama, et al. 2024</u>; <u>Li, et al. 2020</u>). This knowledge gap makes it unclear whether such bee products share the pharmacological properties of the mother plant or constitute distinct chemical resources.

*Epimedium sagittatum* is a renowned traditional Chinese medicinal herb (<u>Liu, et al. 2019</u>). It is characterized by a rich flavonoid content and significant bioactivity (<u>Wang, et al. 2024</u>). Among its major bioactive components, icariin, which is predominantly accumulated in the leaves, plays a crucial role in regulating anti-inflammatory and immune responses (<u>Ji, et al. 2025</u>). *Epimedium* species are also crucial botanical sources for BP production. Yet, research on *Epimedium* bee pollen (EBP) is entirely lacking. To establish EBP as a novel functional ingredient, its chemical profile must be defined not only relative to its source plant but also within the broader context of existing bee pollen products (<u>Liu, et al. 2024</u>; <u>Valverde, et al. 2023</u>). Specifically, its distinctiveness compared to standard commercial pollens remains uncharacterized, and it is completely unknown whether EBP retains the leaf-specific bioactive markers (e.g., icariin) or exhibits a divergent flavonoid pattern.

In this study, we used a UPLC–MS/MS targeted metabolomic platform, and we first performed a horizontal comparison between EBP and widely consumed commercial BPs (BBP and CBP) to benchmark their chemical signatures and pinpoint their unique compositional advantages. More importantly, a vertical comparative analysis was conducted between EBP and the source *Epimedium* plant leaf to quantify the extent of tissue-specific metabolic divergence. By clarifying these multidimensional differences and employing network pharmacology, this research aims to determine whether EBP functions as an independent, high-value functional resource, providing scientific evidence to better understand the distinct chemical characteristics of medicinal plant-derived bee pollen.

## 2. Materials and Methods

### 2.1. Collection and pollen analysis experiment of the BP sample

Multiple pollen samples of *Apis mellifera* were collected from a beekeeping base in Jinhua City, Zhejiang Province, China (longitude: 119.66°, latitude: 29.10°). Honeybees from the same beekeeping base were transported to different sites in Jinhua that cultivate different plant species to collect three BP types (EBP, BBP, and CBP; Figure S1A). All samples were freeze-dried and stored at −80°C until analysis. Three samples of each BP type were selected, yielding a total of nine samples. The samples were air-dried at 30°C and 80% relative humidity, and then dehumidified for 12 h. The pollen samples were homogenized and stored at-− 24°C under argon gas for further analysis. Concurrently, pollen analysis experiments revealed that the pollen from *Epimedium sagittatum*, *Camellia sinensis*, and *Brassica rapa* constituted over 98% of the plant material sources (Figure S2 and Table S1) (<u>Vallese, et al. 2024</u>).

### 2.2. Preparation of BP sample extracts

Weigh 1.000 ± 0.005 g of oven-dried BP sample to constant weight using an electronic balance into a 50 mL screw-cap centrifuge tube. Add 20 mL of 75% ethanol solution, mix thoroughly, and place in an ultrasonic cleaner at 200 W for 20 min in a 50°C water bath. Cool to room temperature, centrifuge at 4000 r/min for 10 min. Repeat the extraction twice. Transfer all supernatant to a 50 mL volumetric flask, dilute to the mark with 75% ethanol solution, and mix thoroughly for analysis of total flavonoids, total polyphenols, and total antioxidants. Prepare the blank solution following the same procedure as the experimental sample.

### 2.3. Determination of total flavonoids in three BPs

#### 2.3.1. Establishment of the rutin standard curve

An aluminum nitrite colorimetric method was used to determine the compounds. Rutin was used as the standard for preparing the standard curve (<u>Han, et al. 2026</u>). An accurately weighed amount of 9.48 mg of rutin, dried to constant weight, was placed into a 50 mL volumetric flask, to which 75% ethanol was added to bring the volume to 50 mL. Volumes of 0.4, 1.2, 2.4, 2.8, and 3.2 mL were aliquoted into a light-protected volumetric flask and treated with 0.4 mL of 5% sodium nitrite. The mixture was shaken thoroughly and allowed to stand for 6 min. Subsequently, 0.4 mL of 10% aluminum nitrate solution was added, followed by thorough shaking, and the mixture was allowed to stand for an additional 6 min. Next, 4 mL of a 4.3% NaOH solution was added, and water was added to the mark. After allowing the solution to stand for 15 min, a blank reference prepared without rutin was used to measure absorbance at 510 nm using a UV-Vis Spectrophotometer (Model MV-1800; Shimadzu Corporation, Kyoto, Japan) (<u>Li, et al. 2012</u>).

#### 2.3.2. Determination of total flavonoid content

Using a 10 mL light-protected colorimetric tube, 1 mL of the extract was combined with 0.4 mL of 5% sodium nitrite. The mixture was shaken thoroughly and allowed to stand for 6 min. Subsequently, 0.4 mL of 10% aluminum nitrate solution was added, and the mixture was shaken again, then allowed to stand for an additional 6 min. Then, 4 mL of 4.3% NaOH was added, and the volume was adjusted to the mark with water, followed by a 15-minute standing period. The absorbance was measured at 510 nm using a UV-visible spectrophotometer (MV-1800, Shimadzu Corporation), and the flavonoid content was calculated using a standard curve.

### 2.4. Determination of total polyphenols in three types of BP

#### 2.4.1. Establishment of a calibration curve

Standard stock solutions of 0.25, 0.5, 1.0, 1.5, 2.0, 2.5, 3.0, and 3.5 mL were added to gallic acid in separate 50 mL volumetric flasks. Water was added to the mark to prepare standard solutions at concentrations of 0.025, 0.05, 0.10, 0.15, 0.20, 0.25, 0.30, and 0.35 mg/mL. Then, 0.2 mL of each concentration of the gallic acid standard solution was transferred into a 10 mL colorimetric tube. Phloroglucinol reagent (0.5 mL) was added to each tube, shaken well, and allowed to stand for 3 min. Then, 1.5 mL of 10.0% sodium carbonate solution was added to each tube, the volume was adjusted to the mark with water, and the tubes were allowed to stand at room temperature in the dark for 20 min. Ethanol (75%) was used as the blank control. Absorbance was measured at 765 nm using a spectrophotometer. Finally, a standard curve and a linear equation were plotted using the mass concentrations (mg/mL) of gallic acid solutions (0.025-0.35 mg/mL) on the x-axis and absorbance on the y-axis (<u>Raposo, et al. 2024</u>).

#### 2.4.2. Determination of total polyphenol content

Sample (0.2 mL) and blank solution (0.2 mL) were pipetted into separate 10 mL colorimetric tubes. Then, 0.5 mL of phloroglucinol reagent was added, thoroughly mixed, and allowed to stand for 3 min. Then, 10.0% sodium carbonate solution (1.5 mL) was added, mixed thoroughly, diluted with water to the mark, and allowed to stand at room temperature in the dark for 20 min. The absorbance of the sample solution was measured at 765 nm using the blank sample solution as a reference, and the concentration of total polyphenols in the test sample solution was calculated using the standard curve.

### 2.5. Determination of total antioxidant capacity in BP

#### 2.5.1. Preparation of test specimens

The extract was taken from the BP sample prepared in Section 2.2. To establish a standard curve, the standard was diluted with FeSO□ diluent to create a series of standard solutions at various concentrations (0, 0.15, 0.3, 0.6, 0.9, 1.2, and 1.5 mmol/L Trolox) (<u>Munteanu and Apetrei 2021</u>).

#### 2.5.2. Determination of antioxidant properties of samples

A 1.5 mol/L FeSO□ standard solution was prepared, followed by the creation of solutions of 1.2, 0.9, 0.6, 0.3, and 0.15 mol/L FeSO□. Take 180 μL of FRAP working solution (prepare fresh for immediate use) and mix thoroughly with 20 μL of BP sample solution. Incubate at 37°C for 5 min, then measure the absorbance at 593 nm. The total antioxidant capacity of the BP sample is expressed as FeSO□ equivalents per gram of BP, with units of mmol/g.

#### 2.5.3. Data processing

A standard curve was plotted using the concentration of the standard samples (x-axis, mmol/L Trolox) as the horizontal coordinate and the corresponding absorbance values (y-axis) as the vertical coordinate (y = 0.6721x + 0.0498, R² = 0.9991). The measured absorbance values for the BP samples were substituted into the standard curve equation to calculate the sample solution’s antioxidant capacity, expressed as mmol/L Trolox equivalents. The total antioxidant capacity (μmol Trolox/g fresh weight) was calculated using the following formula:

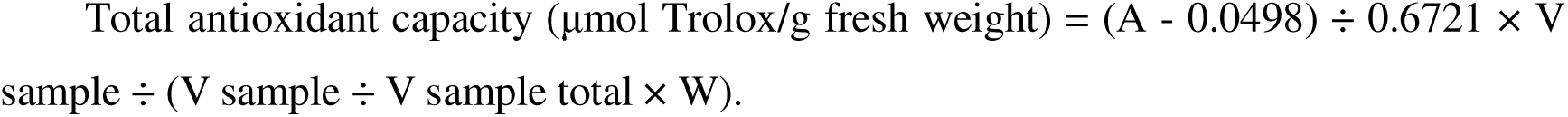

### 2.6. UPLC-MS/MS experiments of BP extract

#### 2.6.1. Preparation of BP extract

Samples were prepared and extracted using methods provided by Metware Biotechnology Co., Ltd. (Wuhan, China). BP samples were placed in a freeze dryer (Scientz-100F) and vacuum freeze-dried for 63 h. Subsequently, an MM 400 grinder (Retsch, Haan, Germany) was used to grind the samples at 30 Hz for 1.5 min, producing a powder. From this powdered material, 30 mg of the sample powder was weighed using an electronic balance and then mixed with 1500 μL of a 70% methanol–water internal standard extraction solution, which had been pre-cooled to −20°C. If less than 30 mg was obtained, the extraction solvent volume was adjusted accordingly to 1,500 µL per 30 mg of sample. The internal standard extraction solution was prepared by dissolving 1 mg of the standard substance in 1 mL of 70% methanol–water, yielding a stock solution of 1,000 μg/mL. This stock solution was further diluted with 70% methanol (Merck, Darmstadt, Germany) to prepare the internal standard solution at 250 μg/mL. The samples were vortexed every 30 min for 30 s, for a total of 6 vortexing sessions, then centrifuged at 12,000 rpm for 3 min. The supernatant was then collected and filtered through a microporous membrane (0.22 μm pore size) to obtain the sample solution, which was subsequently stored in a sample vial for further analysis of active components in pollen by ultra-performance liquid chromatography (UPLC)-tandem mass spectrometry (MS/MS) (ExionLC™AD, https://sciex.com.cn/).

#### 2.6.2. UPLC and MS conditions

Metabolite detection was performed on an ExionLC™ AD ultra-performance liquid chromatography system coupled to a triple quadrupole-linear ion trap mass spectrometer equipped with an electrospray ionization (ESI) source. Chromatographic separation was achieved using an Agilent SB-C18 column (1.8 μm, 2.1 mm × 100 mm). The mobile phase consisted of water with 0.1% formic acid (phase A) and acetonitrile with 0.1% formic acid (phase B). The gradient program was as follows: 5% B at 0 min, linearly increased to 95% B within 9.0 min, held at 95% B for 1.0 min, then returned to 5% B and equilibrated until 14.0 min. The flow rate was 0.35 mL/min, the injection volume was 2 μL, and the column temperature was maintained at 40□.

The mass spectrometer was operated in both positive and negative ion modes under multiple reaction monitoring (MRM) conditions. The ESI source parameters were set as follows: ion source temperature, 500□; ion spray voltage, +5500 V in positive mode and −4500 V in negative mode; gas I, 50 psi; gas II, 60 psi; curtain gas, 25 psi; and collision gas, medium. Declustering potential and collision energy were optimized for individual MRM ion pairs. Peak detection, integration, and correction across samples were performed based on the uniform retention time and ion-pair information for each metabolite (<u>Fraga, et al. 2010</u>).

To monitor analytical stability, a pooled quality control (QC) sample was prepared by mixing equal aliquots of all sample extracts and was injected repeatedly throughout the analytical sequence. The reproducibility of retention time and signal intensity in QC runs was used to assess instrument stability. Only metabolites showing consistent peak recognition and acceptable signal reproducibility across QC samples were retained for downstream analysis. Raw peak areas were exported after peak integration and alignment for subsequent statistical analyses.

### 2.7. Description of statistical analysis

#### 2.7.1. Statistical analysis

All biochemical measurements were performed using three biological replicates (n = 3), and the results are presented as mean ± standard deviation (SD). Statistical analyses for total flavonoids, total polyphenols, and antioxidant capacity were performed using IBM SPSS Statistics 26.0 (IBM Corp., Armonk, NY, USA). Differences among the three bee pollen types were evaluated using one-way analysis of variance (ANOVA), followed by Tukey’s honestly significant difference (HSD) test for multiple comparisons. A p-value < 0.05 was considered statistically significant.

Multivariate statistical analyses of metabolomic data were performed in R 4.5.2 (R Foundation for Statistical Computing, Vienna, Austria). Unless otherwise stated, metabolomic data visualization and downstream multivariate analyses were based on normalized peak-area matrices exported from the UPLC–MS/MS platform.

#### 2.7.2. Principal component analysis (PCA)

Principal component analysis (PCA) was used as an unsupervised method to assess the overall variation in metabolite composition among samples. Prior to PCA, the metabolite peak-area matrix was subjected to data preprocessing in the following order: peak-area extraction, sample-wise matrix assembly, normalization, log_2_ transformation where appropriate, and unit-variance scaling. PCA was then performed using the prcomp function (www.r-project.org) in R. Score plots were used to visualize sample separation among EBP, BBP, and CBP (Figure S3, Table S2).

#### 2.7.3. Hierarchical Cluster Analysis and Pearson Correlation Coefficients

Hierarchical cluster analysis (HCA) and Pearson correlation coefficient (PCC) analysis were conducted to evaluate the similarity among samples and the clustering pattern of metabolites. Analyses were performed in R using the ComplexHeatmap package. The input matrix consisted of normalized metabolite signal intensities. For HCA, unit-variance-scaled data were visualized as heatmaps with dendrograms. For PCC analysis, pairwise Pearson correlation coefficients were calculated across biological replicates using the normalized metabolite matrix, and the resulting correlation matrix was displayed as a heatmap.

#### 2.7.4. Differential metabolites selected

To identify metabolites contributing to pairwise differences between EBP and the two reference bee pollens, supervised orthogonal partial least squares-discriminant analysis (OPLS-DA) was performed using the MetaboAnalystR package in R. Before model construction, metabolite intensity data were log_2_-transformed and mean-centered (Figure S5, Table S2). Variable importance in projection (VIP) values were extracted from the OPLS-DA model, and differential metabolites were defined using the combined criteria of VIP > 1 and absolute log2 fold change (|log_2_FC|) ≥ 1.0.

To reduce the risk of model overfitting, permutation testing (200 permutations) was conducted for each OPLS-DA model. The resulting differential metabolites were subsequently used for metabolite classification, overlap analysis, and pathway annotation. In the present study, OPLS-DA was used as an auxiliary discriminative tool for metabolite prioritization rather than as a standalone basis for biological interpretation.

#### 2.7.5. KEGG annotation and enrichment analysis

Differential metabolites identified in pairwise comparisons were annotated against the KEGG Compound database (http://www.kegg.jp/kegg/compound/) and mapped to the KEGG Pathway database (http://www.kegg.jp/kegg/pathway.html) for functional classification. KEGG enrichment analysis was performed to assess whether specific metabolic pathways were overrepresented among the differentially abundant metabolites. In this study, KEGG results were used to support pathway-level interpretation of compositional differences, rather than to infer direct biological mechanisms or causal metabolic regulation.

Identified metabolites were annotated using the KEGG Compound database (http://www.kegg.jp/kegg/compound/), and annotated metabolites were then mapped to the KEGG Pathway database (http://www.kegg.jp/kegg/pathway.html).

#### 2.7.6. Network pharmacology analysis

To further prioritize metabolites of potential interest within the shared metabolite set, an auxiliary network pharmacology analysis was performed as an *in silico* screening step. This analysis was not intended to validate therapeutic efficacy, but rather to identify metabolites that may deserve further attention based on commonly used activity-related filters.

Candidate compounds were first screened in the TCMSP database using oral bioavailability (OB) ≥ 30% and drug-likeness (DL) ≥ 0.18. For compounds not covered by TCMSP (https://old.tcmsp-e.com/tcmsp.ph), pharmacokinetic and drug-likeness properties were evaluated using SwissADME (http://www.swissadme.ch/). Compounds with high gastrointestinal absorption and at least two positive drug-likeness criteria among Lipinski, Ghose, Veber, Egan, and Muegge were retained. Potential targets of the retained compounds were then collected from TCMSP, SwissTargetPrediction (http://www.swisstargetprediction.ch/), and SymMap (https://www.symmap.org/), and target names were standardized using UniProt Gene Symbols.

Disease-related targets were retrieved from DisGeNET (https://www.disgenet.org/), SymMap, and TTD. Overlapping targets between compound-associated targets and disease-associated targets were identified and used to construct component–target–disease associations. Protein–protein interaction (PPI) analysis of overlapping targets was further conducted using the STRING database (http://cn.string-db.org/). In the present study, the purpose of this analysis was to support activity-related prioritization of metabolites associated with flavonoids highlighted by metabolomic analysis, rather than to provide independent evidence of health functionality.

## 3. Results and Discussion

### 3.1. Biochemical and metabolomic characteristics of EBP

Biochemical analysis revealed significant differences among the three bee pollen types from different botanical origins. EBP exhibited the highest total flavonoid content (4.75 mg/g), significantly exceeding that of both BBP and CBP (*p* < 0.05; Figure S1B), while its total polyphenol content and antioxidant capacity were also significantly higher than those of CBP (*p* < 0.05). These results suggest that EBP was distinguishable from the other two bee pollen types using major biochemical indices. PCA further supported this pattern by showing significant compositional separation among the three bee pollen types (Figure S6; Table 1). Considering the broad commercial application and recognized health-related value of CBP and BBP, the present physicochemical results provide supportive information for the further exploration of EBP, particularly given its relative advantages in total flavonoid content and antioxidant capacity (<u>Lü, et al. 2025</u>; <u>Yuan, et al. 2024</u>). Together, these results indicate that EBP possessed a more distinct chemical composition, which provided the rationale for subsequent differential component analysis.

**Table 1:** Substance classification and quantity of metabolites of three kinds of bee.

| Class | EBP | BBP | CBP |
| --- | --- | --- | --- |
| Alkaloids | 200 | 202 | 181 |
| Flavonoids | 330 | 362 | 358 |
| Lignans and Coumarins | 52 | 51 | 58 |
| Phenolic acids | 165 | 161 | 180 |
| Quinones | 14 | 16 | 14 |
| Steroids | 2 | 2 | 1 |
| Tannins | 9 | 7 | 34 |
| Terpenoids | 170 | 167 | 158 |
| Others | 131 | 128 | 132 |
| Total | 1073 | 1096 | 1116 |

pollen.

#### 3.1.1. Unique metabolite features of EBP relative to BBP and CBP

To better characterize the unshared compositional features of the three bee pollen types, unique metabolites were evaluated to delineate their sample-specific chemical characteristics (<u>Ricigliano, et al. 2022</u>; <u>Zuccato, et al. 2017</u>). Comparative metabolomic analysis revealed 19, 14, and 271 unique metabolites in EBP, BBP, and CBP, respectively (Figure 1A, Table S3), indicating divergence in their secondary metabolite profiles. Notably, although EBP did not have the highest number of unique metabolites, it exhibited a highly concentrated, distinct metabolite profile. In contrast, CBP was characterized by a broad spectrum of unique components, whereas BBP displayed the least diverse exclusive chemical profile.

**Figure 1.**
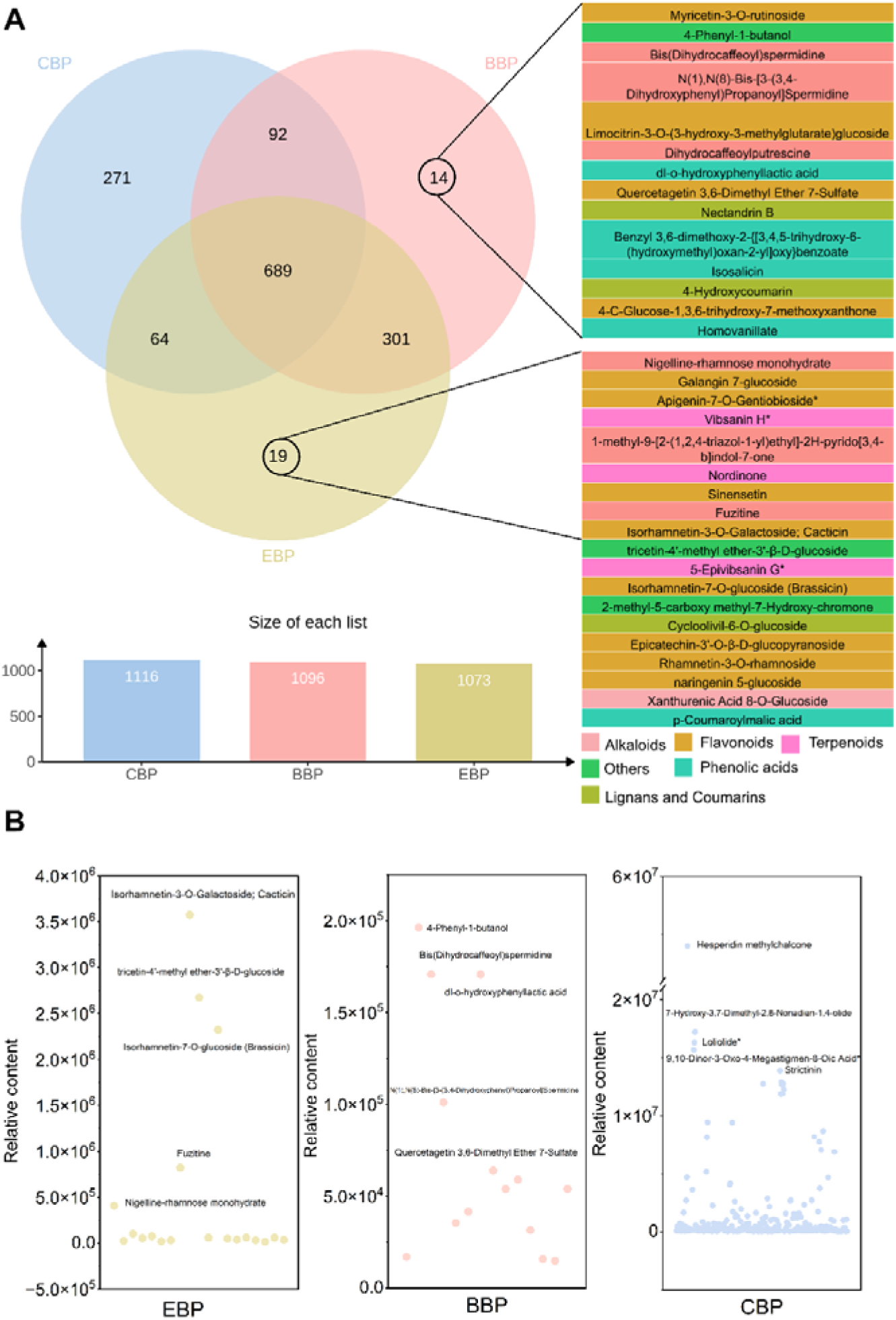
(A) Venn diagram of total metabolites of three types of BP. (B) Changes in the relative content of pollen characteristic metabolites in EBP (a), BBP (b), and CBP (c). CBP: Camellia sinensis bee pollen; BBP: Brassica rapa bee pollen; EBP: *Epimedium* bee pollen. For the data shown in Panel B, different letters indicate significant differences at p < 0.05. Data are represented as the mean ± standard error of the mean SEM (n=3).

Chemical classification revealed that the majority of these unique metabolites were flavonoids and their derivatives (Table 2), highlighting this class as a crucial dimension driving the chemical divergence among the three bee pollen types. Previous studies also indicated that flavonoids are the principal bioactive compounds in some BP types (<u>Lv, et al. 2015</u>). While flavonoids are common in plants and bee pollen, their specific distribution here suggests that the differences in unique metabolite profiles may be associated, at least in part, with differences in the botanical origins of each bee pollen type (<u>Denisow and Denisow-Pietrzyk 2016</u>). When evaluating relative abundances, EBP showed a clear concentration of specific compounds. Among its 19 unique metabolites, several were highly concentrated, particularly isorhamnetin-3-O-galactoside (cacticin), tricetin-4 ′ -methyl ether-3 ′ - β-D-glucoside, and isorhamnetin-7-O-glucoside (brassicin), all with relative peak areas exceeding 2.0 × 10^6^ (Figure 1B-a). On the other hand, the unique metabolites in BBP were few and present at much lower levels, mostly peaking below 2.0 × 10^5^ (Figure 1B-b). By comparison, CBP had the largest number of unique metabolites and the highest signal intensities (reaching 107, Figure 1B-c). However, instead of being concentrated in a single chemical class, the unique compounds in CBP were scattered across a wide range of chemical types.

**Table 2:** Classification of characteristic metabolites and metabolite quantities of three bee pollen.

| Class | EBP | BBP | CBP |
| --- | --- | --- | --- |
| Alkaloids | 4 | 3 | 13 |
| Flavonoids | 8 | 4 | 101 |
| Lignans and Coumarins | 1 | 2 | 19 |
| Phenolic acids | 1 | 4 | 52 |
| Quinones | 0 | 0 | 0 |
| Steroids | 0 | 0 | 1 |
| Tannins | 0 | 0 | 26 |
| Terpenoids | 3 | 0 | 36 |
| Others | 2 | 1 | 23 |
| Total | 19 | 14 | 271 |

In summary, the unshared metabolite profiling reveals a highly specialized chemical signature unique to EBP. Unlike RBP and CBP, EBP is specifically enriched for a group of highly abundant flavonoids. This concentrated compositional fingerprint not only definitively establishes the distinct chemical identity of EBP but also highlights its unique phytochemical value.

#### 3.1.2. Differential metabolites associated with EBP

To comprehensively elucidate the chemical signature of EBP, differential metabolite analysis was conducted. While unique metabolite profiling successfully identified the unshared chemical fraction, analyzing differentially accumulated metabolites (DAMs) captures the broader, quantitative metabolic shifts that drive the distinct phenotype of EBP compared to BBP and CBP (<u>Yue, et al. 2025</u>).

Based on multivariate analyses (PCA and OPLS-DA; Figure S5C-D), 449 (vs. BBP) and 1,085 (vs. CBP) differential metabolites were identified in EBP relative to BBP and CBP, respectively (Figure 2A, B; Table S4), indicating marked metabolic divergence (<u>Liu, et al. 2025</u>). In both comparisons, flavonoid compounds represented the dominant class of discriminative metabolites (Figure 2D, E), suggesting that the chemical differences associated with EBP were concentrated to a considerable extent in flavonoid and related constituents. This result is also consistent with previous biochemical analyses and with the identification of unique metabolites.

**Figure 2.**
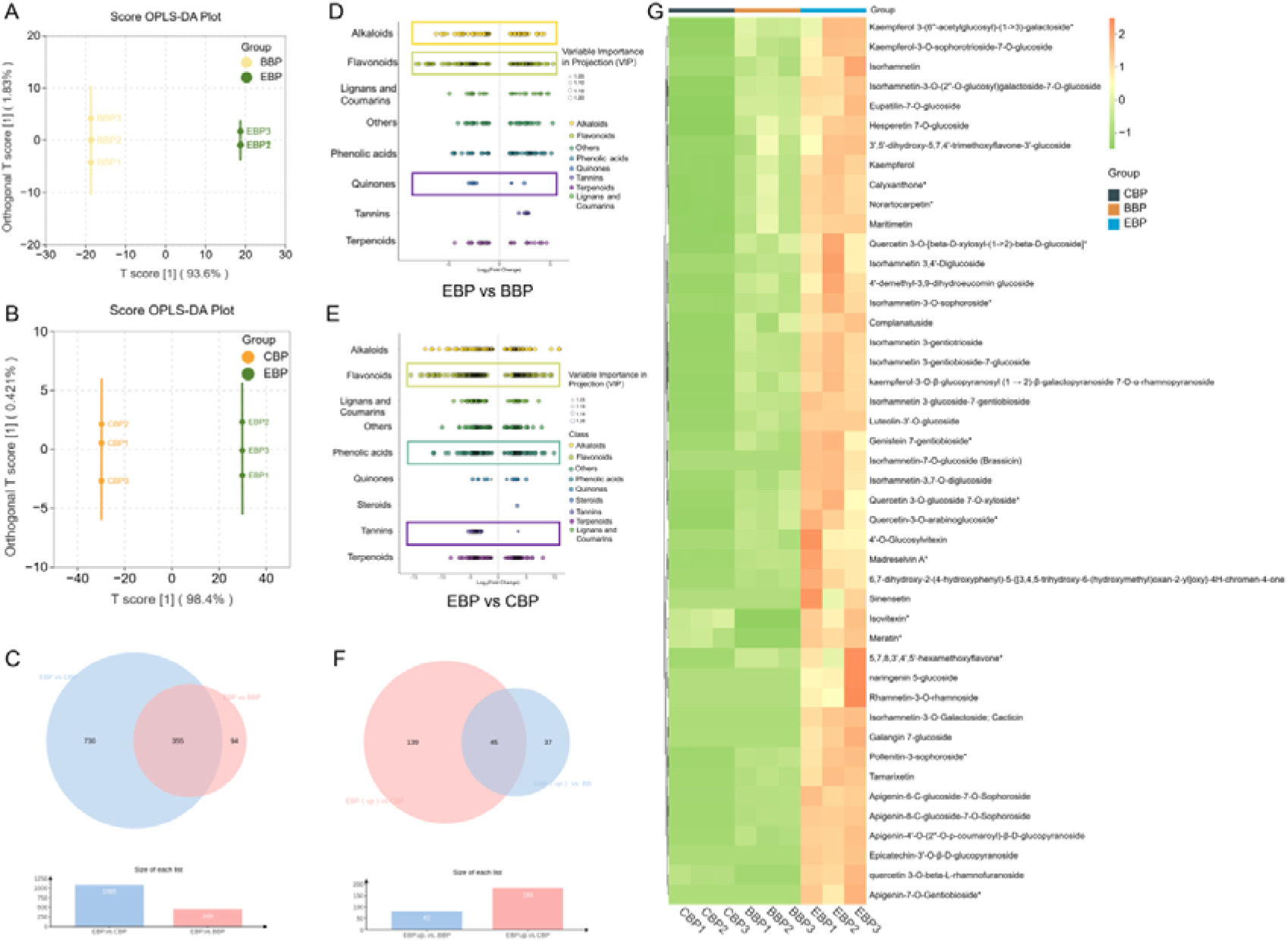
OPLS-DA score chart of (A) EBP vs BBP group and (B) EBP vs CBP group. (C) Venn diagram of significantly different metabolites between EBP and the other two BPs; Scatter plot of differential metabolites in (D) EBP vs BBP group and (E) EBP vs CBP group. (F) Venn diagram showing significantly higher levels of flavonoid metabolites in EBP than in BBP and CBP. (G) 45 flavonoids were upregulated considerably in EBP. CBP: *Camellia sinensis* bee pollen; BBP: *Brassica rapa* bee pollen; EBP: *Epimedium* bee pollen. Different letters indicate significant differences at p < 0.05. Data are represented as the mean ± standard error of the mean (SEM).

More importantly, the differential flavonoids in EBP showed a certain degree of compositional relatedness. Among the 355 shared DAMs detected in both pairwise comparisons, 45 flavonoids were consistently upregulated in EBP (Figure 2C, F, G). Several representative compounds, including kaempferol, tamarixetin, sinensetin, norartocarpetin, and isorhamnetin-3-O-galactoside, belong to structurally related flavonoid-associated metabolites, and some of them share common modification patterns, such as oxygenation, methylation, or glycosylation (<u>Shen, et al. 2022</u>; <u>Zhao, et al. 2018</u>). This suggests that the flavonoid differences in EBP were not simply due to an indiscriminate increase in flavonoid diversity, but rather to the enrichment of a group of related flavonoid metabolites. This metabolic reshaping is robustly supported by KEGG pathway analysis, which demonstrates significant enrichment of the flavonoid biosynthesis pathway (ko00941) in EBP compared to both BBP and CBP (Figure S7) (<u>Wang, et al. 2018</u>; <u>Zhao, et al. 2019</u>). Although pathway enrichment does not constitute direct mechanistic validation, it supports the view that the flavonoid metabolic profile differences associated with EBP were a systematic change.

At the individual metabolite level, kaempferol, tamarixetin, and sinensetin showed relatively large fold changes, whereas norartocarpetin and isorhamnetin-3-O-galactoside were notable for their comparatively high abundance (Figure 3; Table 3). This indicates that the distinctiveness of EBP was supported by both the marked relative increases in some flavonoid metabolites and the substantial accumulation of others, especially flavonoids concentrated.

**Figure 3.**
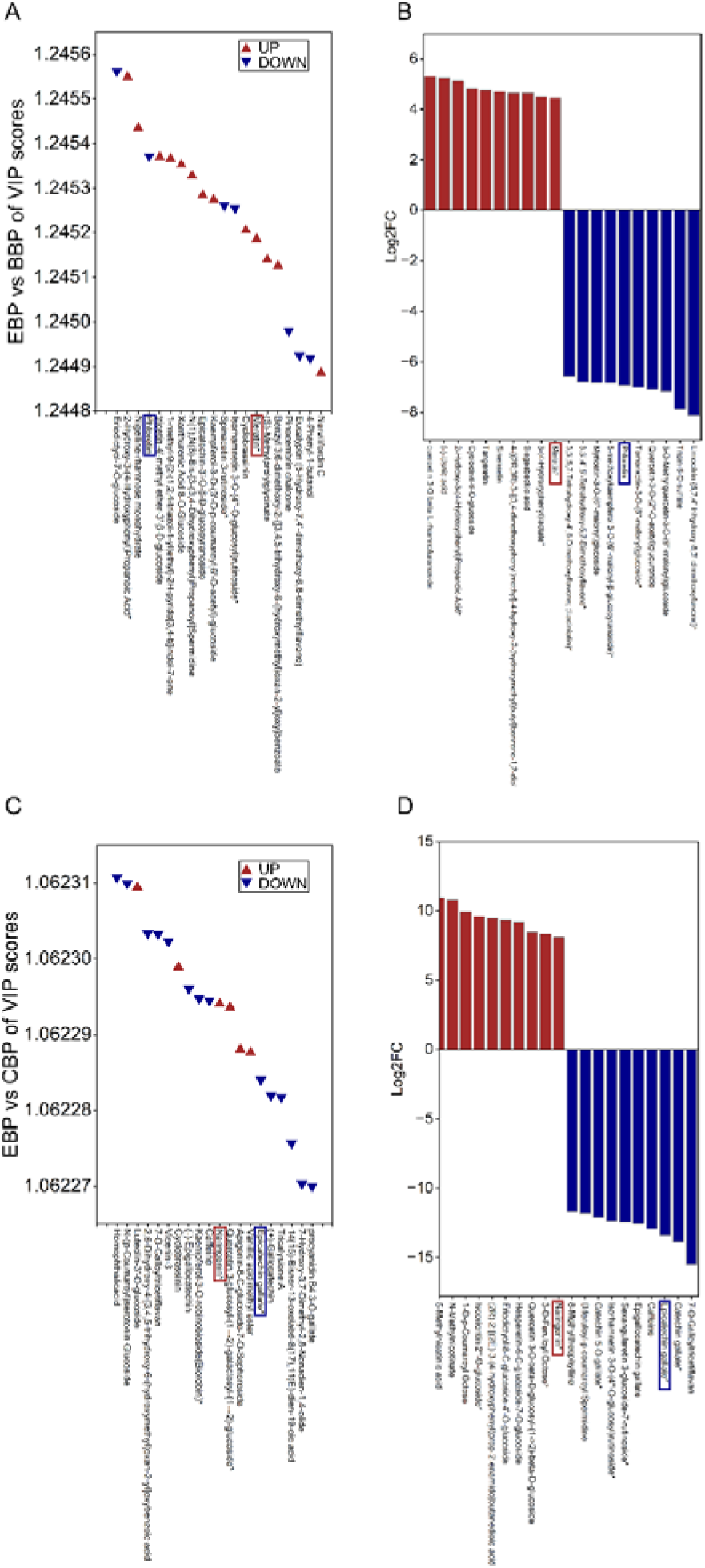
EBP vs CBP of differential metabolite Variable Importance in Projection (VIP) value chart (A) and bar chart of the top 10 substances with differential changes in dif-ferential metabolites (C). EBP vs BBP of differential metabolite VIP value chart (B) and bar chart of the top 10 substances with differential changes in differential metabolites (D). CBP: *Camellia sinensis* bee pollen; BBP: *Brassica rapa* bee pollen; EBP: *Epimedium* bee pollen.

**Table 3:**
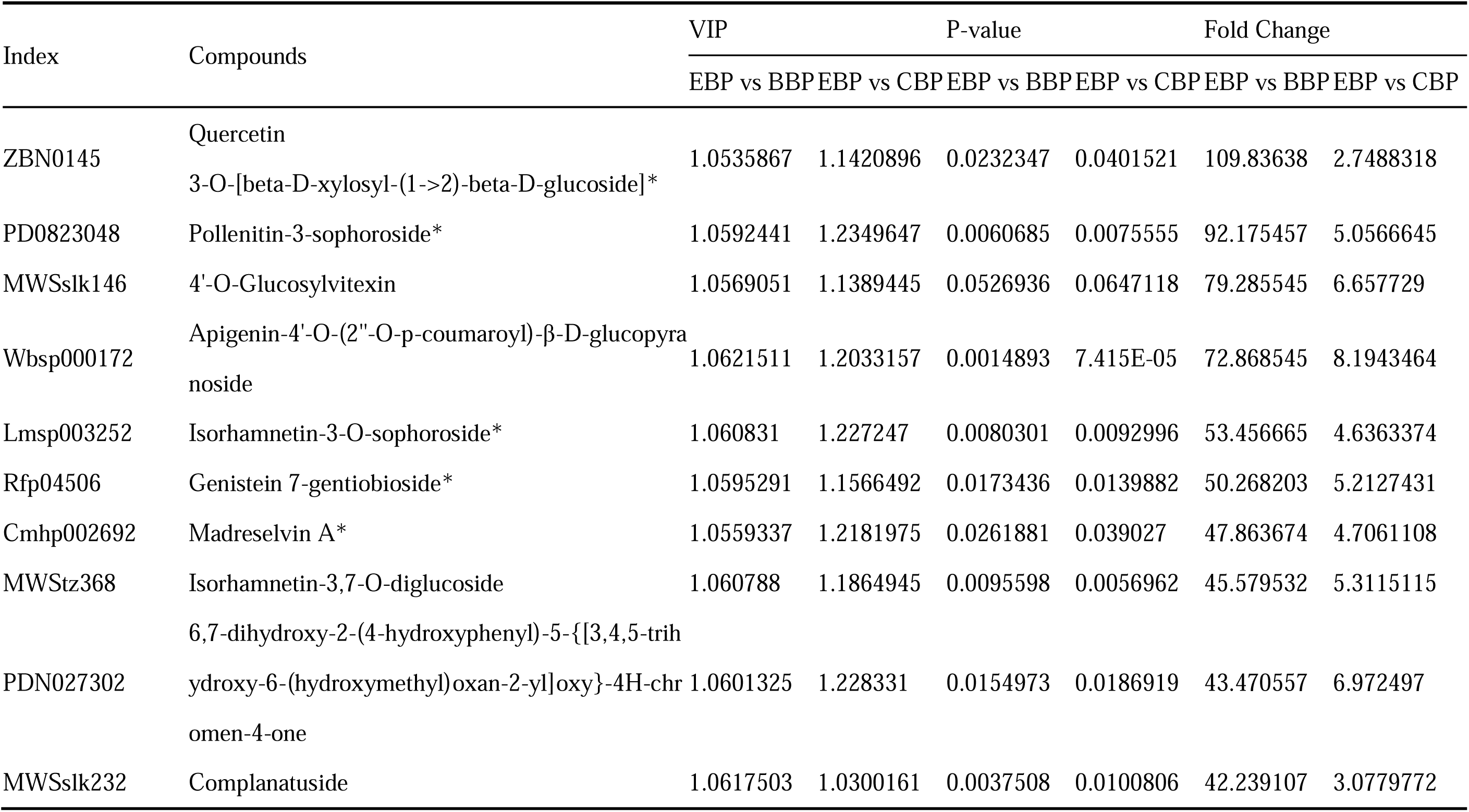
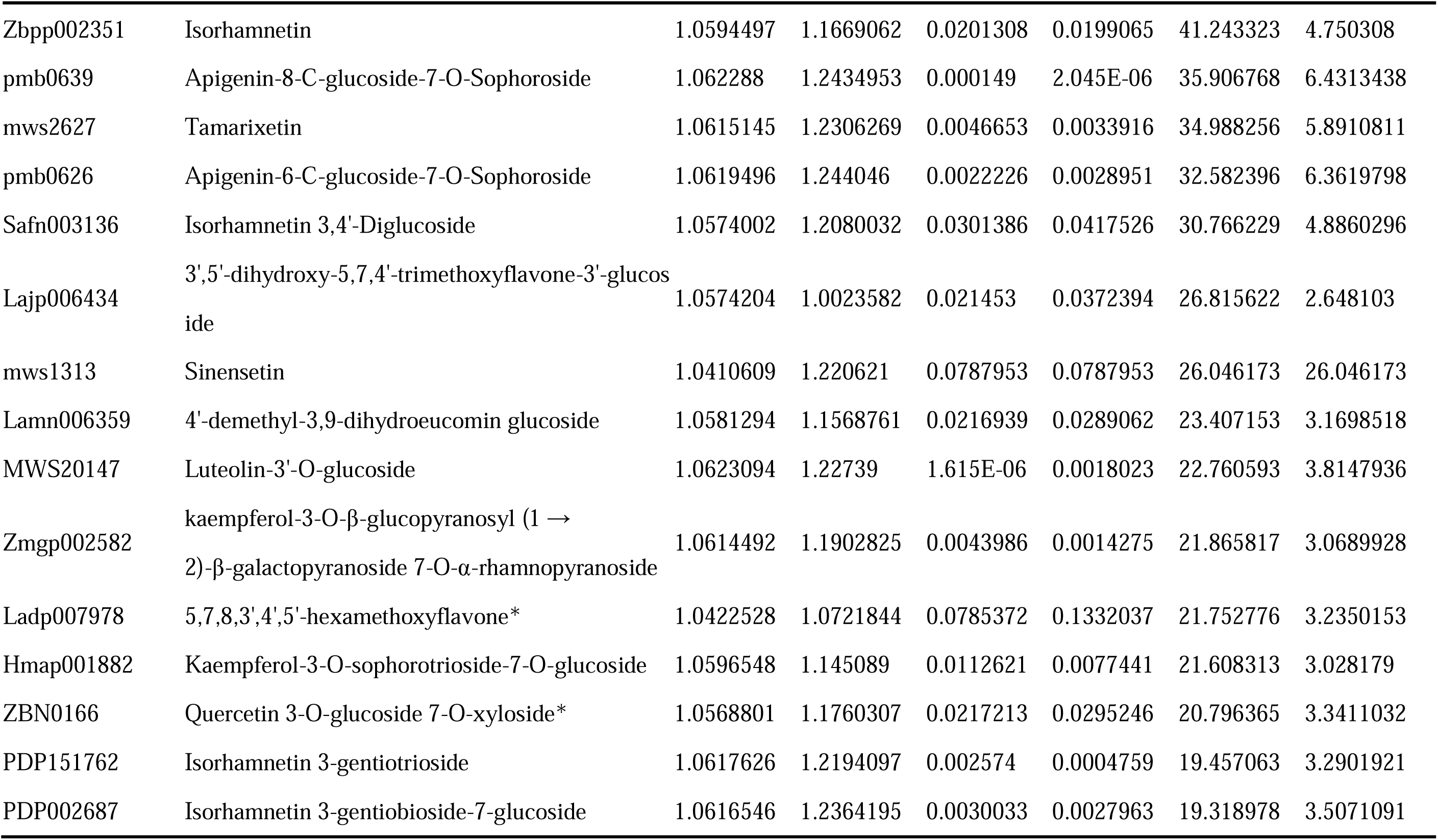
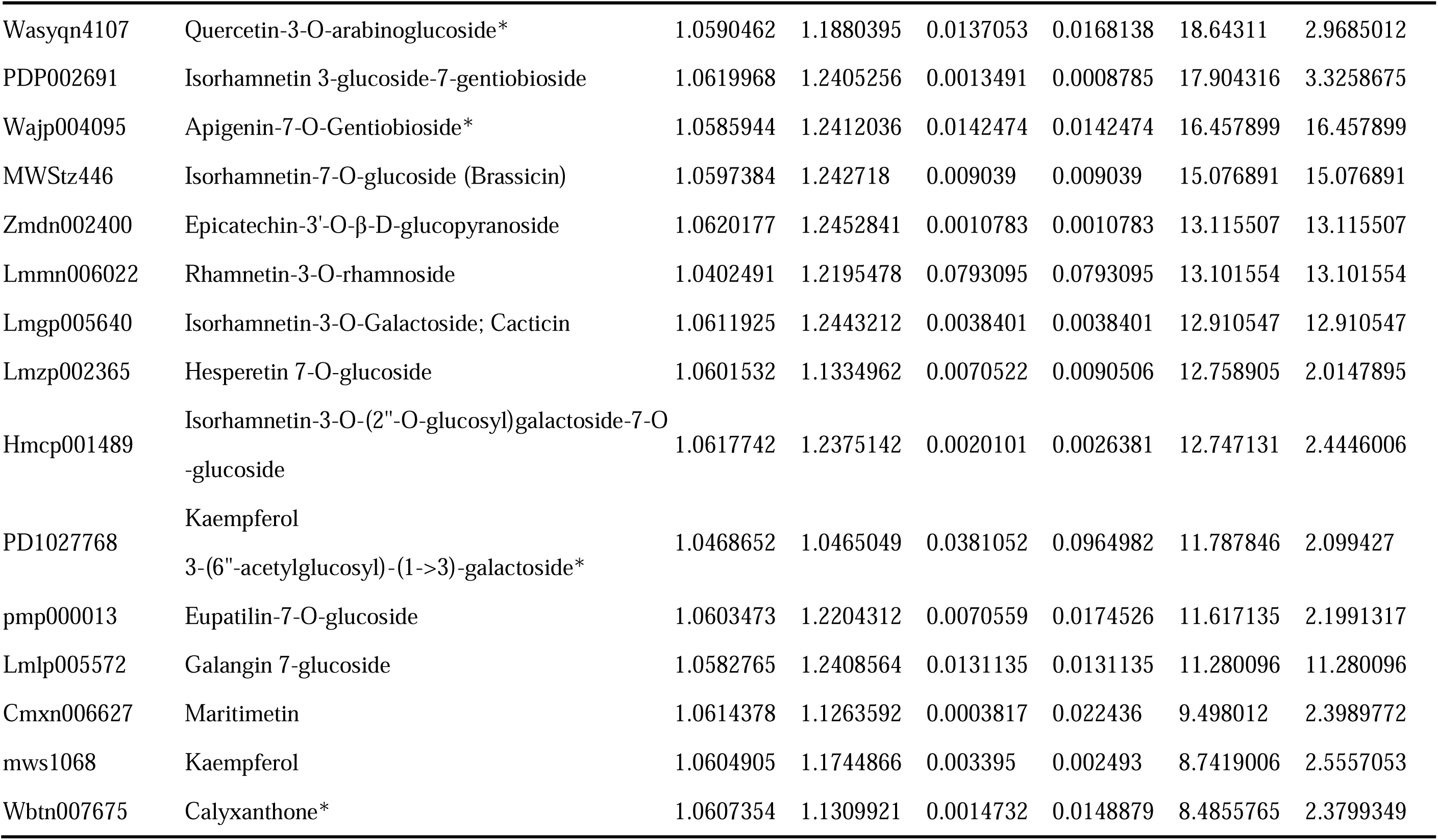
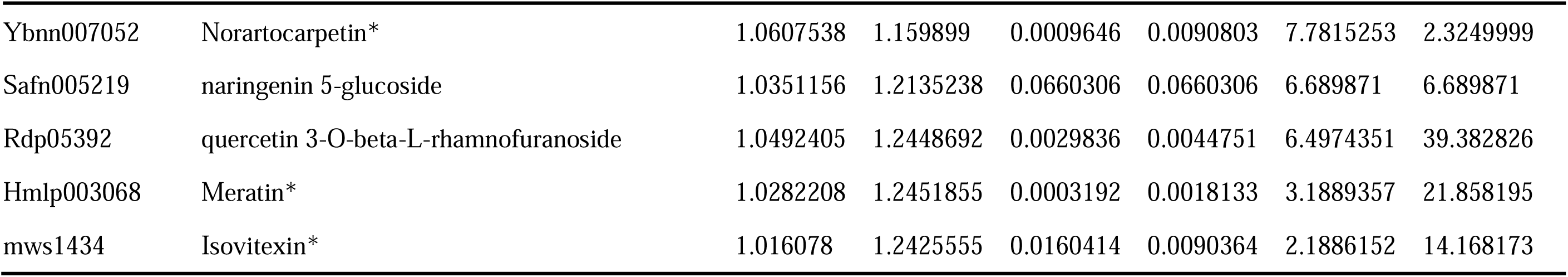
Significantly higher flavonoid metabolites in EBP compared to BBP and CBP.

Collectively, these findings demonstrate that the overarching chemical feature of EBP is the systematic enrichment of a panel of structurally related flavonoids. This highly specialized metabolic profile provides a comprehensive and distinctive chemical basis for the botanical authentication and further utilization of EBP.

### 3.2. Network pharmacology highlights key flavonoid-related metabolites in EBP

To determine whether the chemical distinctiveness of EBP translates into biologically meaningful features, network pharmacology analysis was employed to prioritize the 211 shared secondary metabolites across the three pollen types. Given that directly inferring specific therapeutic effects ultimately depends on actual *in vivo* concentrations and bioavailability, this *in silico* approach was used as a screening tool to assess whether the previously observed flavonoid variations are the most relevant aspect of EBP’s compositional divergence (<u>Shawky 2019</u>). Applying standard ADMET criteria (oral bioavailability ≥ 30%, drug-likeness ≥ 0.18, and probability > 0.12), 25 highly active candidate metabolites were prioritized (Figure S4A, B; Table S5). Notably, EBP retained a broader repertoire of these prioritized constituents (22 metabolites, including 15 flavonoids) compared to BBP (20) and CBP (19) (Figure 4E). This result robustly validates that the functionally relevant potential of EBP is tightly anchored to its flavonoid-enriched profile, solidifying flavonoids as the most noteworthy chemical differentiators (<u>Denisow and Denisow□Pietrzyk 2016</u>).

**Figure 4.**
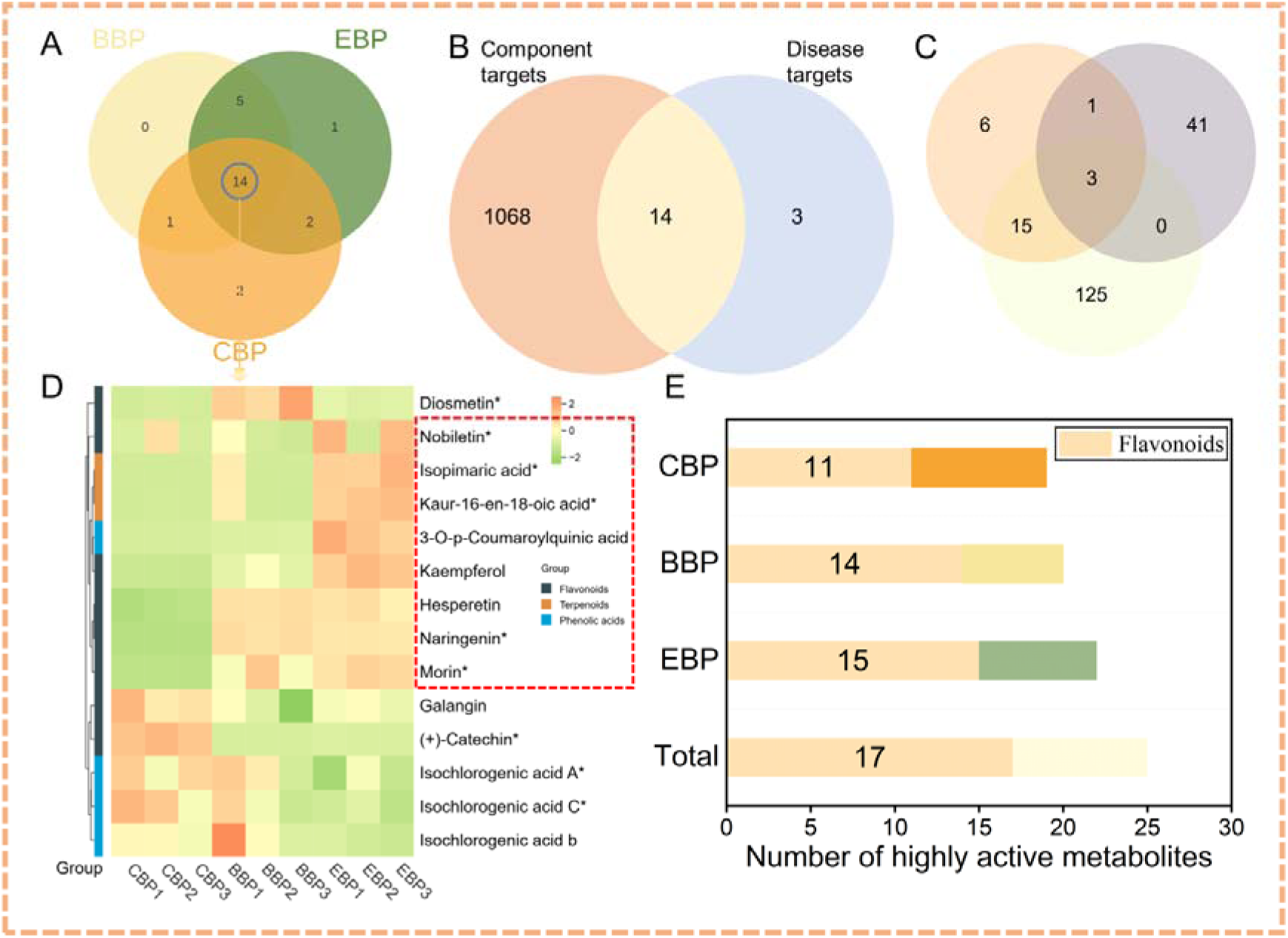
(A) Venn diagram showing the overlap between the three types of highly active substances in pollen. (B) Venn diagram depicting the shared and unique differentially accumulated metabolites between pairs of component targets and disease targets. (C) Venn diagram showing the cross-analysis of metabolites screened by TCMSP, 45 flavonoids upregulated in EBP relative to the other two groups of BP, and metabolites corresponding to disease targets. (D) Heat map of 14 metabolites overlapping with three types of highly active substances in BP. (E) Comparison of the number of highly active metabolites in three types of BP. BP, bee pollen; CBP: *Camellia sinensis* bee pollen; BBP: *Brassica rapa* bee pollen; EBP: *Epimedium* bee pollen.

Strikingly, the distribution of these computationally prioritized metabolites perfectly mirrored the targeted flavonoid enrichment observed in the preceding metabolomic analysis. Among the 14 core candidate metabolites shared across all three pollen types (Figure 4A), eight accumulated at notably higher levels in EBP than in BBP and CBP (Figure 4D; Table 4). Crucially, this dominant cluster is composed entirely of flavonoid derivatives, highlighted by the prominence of morin, kaempferol, and nobiletin, as well as elevated levels of tamarixetin, quercetin, and sinensetin. The high health value of these substances underscores the significance of flavonoids in EBP (<u>Bangar, et al. 2022</u>; <u>Rajput, et al. 2021</u>; <u>Wang, et al. 2021</u>).

**Table 4:**
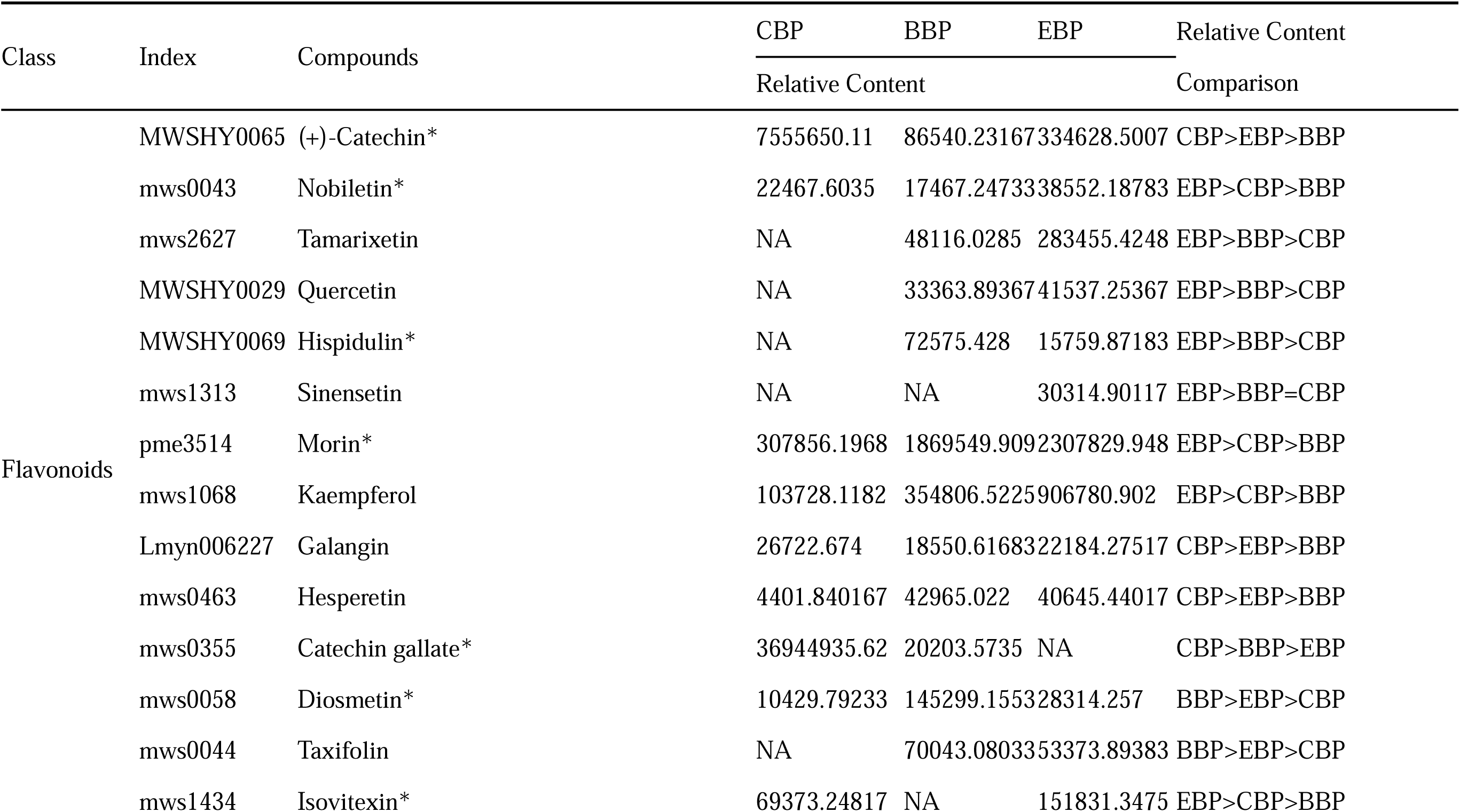
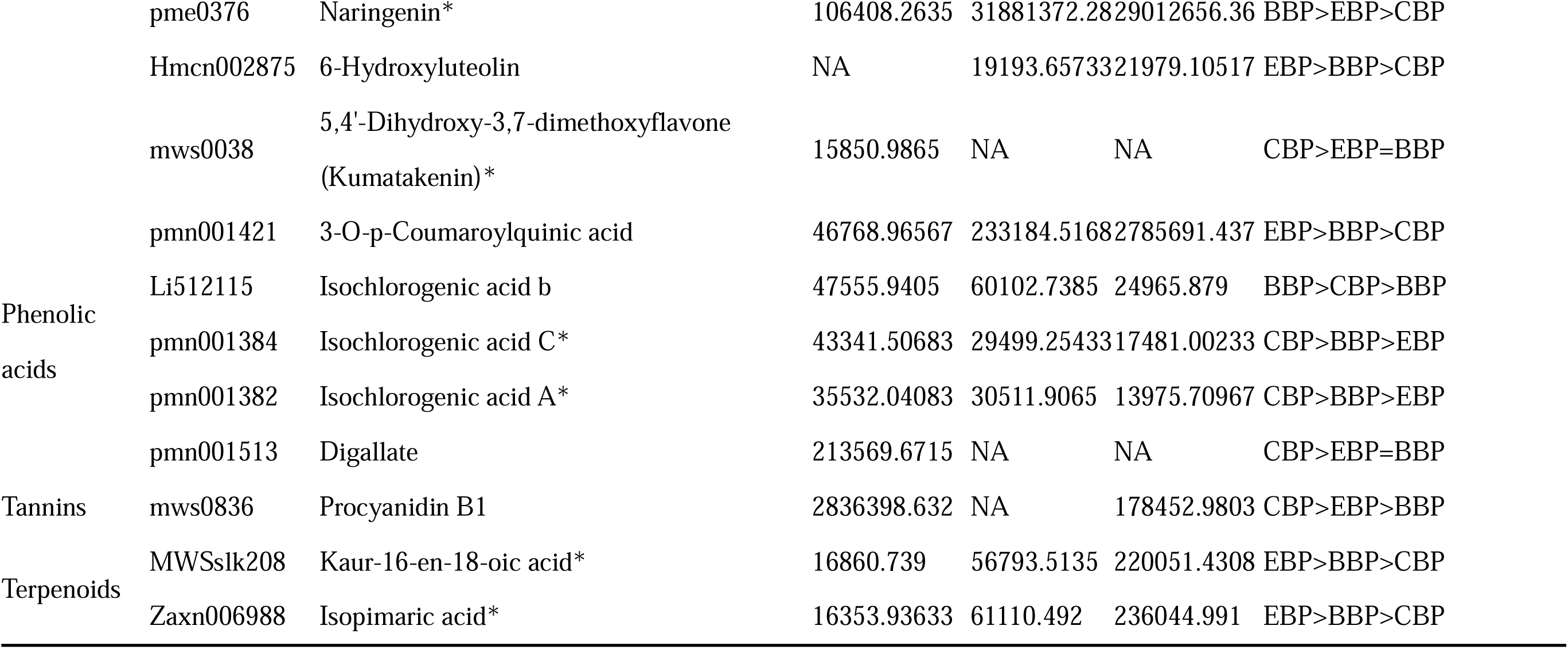
Cross-over as well as content comparison of metabolites screened by the EBP、CBP and CBP with the TCMSP system.

Integrating target predictions via DisGeNET, kaempferol, tamarixetin, and sinensetin emerged as pivotal nodes. Among the 143 metabolites associated with 14 key disease targets, EBP covers 118 (82.5% coverage, Table S6). Among them, kaempferol, tamarixetin, and sinensetin are particularly notable because they simultaneously meet the stringent activity-related criteria and are significantly upregulated in EBP relative to BBP and CBP (Figure 4B, C), especially kaempferol concentration in EBP is 7–8 times higher than in CBP (Figure S8, *p* < 0.05). This convergence of results demonstrates that focusing on these specific flavonoids is not only chemically accurate based on differential analysis but also thoroughly justified from the perspective of activity-related prioritization. In summary, the *in silico* screening provides compelling auxiliary evidence that flavonoids constitute the defining chemical hallmark of EBP.

### 3.3. Distinct metabolite composition of EBP and *Epimedium* leaves

The targeted metabolomic profiling of EBP identified 1,073 secondary metabolites (Figure S6D), indicating a highly complex chemical composition. Compared with the metabolite dataset recently reported for *Epimedium* leaves (<u>Chen, et al. 2024</u>), EBP contained a substantially larger number of detected metabolites, suggesting marked differences in metabolite composition between bee pollen and leaf. Although flavonoids remained the dominant class in both EBP and *Epimedium* leaves (Figure 5A), their compositional features were clearly different. In particular, icariin and its major related derivatives, including epimedin A, epimedin B, and epimedin C, which are characteristic constituents of *Epimedium* leaves, were present at very low levels or were not detected in EBP (Figure 5B, C). By contrast, EBP was characterized by the accumulation of other flavonoid-related metabolites, including several compounds that were not prominent in the leaf dataset. These results indicate that EBP did not simply mirror the metabolite composition of the source plant leaves but instead exhibited a distinct chemical profile.

**Figure 5.**
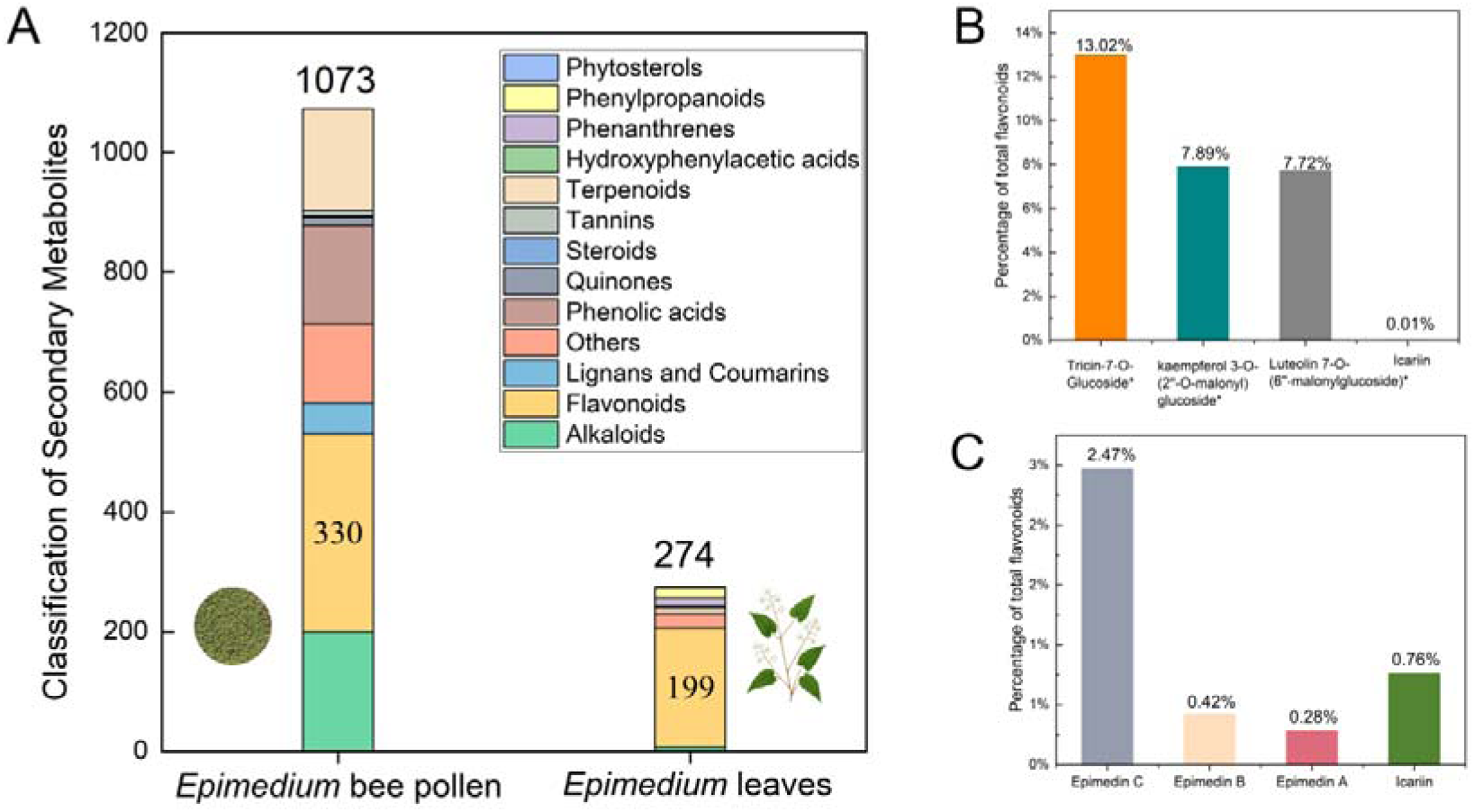
(A) Metabolite classification of EBP and *Epimedium* leaves (<u>Chen, et al. 2024</u>). (B) The three most abundant flavonoids in EBP and the relative content of *icariin*. (C) The flavonoids with the highest relative content in *Epimedium* (<u>Wang, et al. 2023</u>).

Such divergence may be associated with tissue-dependent differences in plant metabolism, since reproductive tissues and vegetative tissues often differ in metabolite accumulation patterns. In addition, because bee pollen is formed after pollen collection and packing by bees, the final metabolite profile of EBP may also be influenced by biochemical transformation during pollen processing, including possible enzymatic modification by bee-derived factors (<u>Isah 2019</u>; <u>Li, et al. 2024a</u>). Although the relative contributions of these processes cannot be resolved here, the present results clearly show that EBP represents a chemically distinct matrix compared with that of *Epimedium* leaves.

A similar divergence between medicinal plant materials and their corresponding bee pollens has also been reported in *Camellia sinensis*, *Lycium barbarum*, and *Schisandra chinensis* (<u>An, et al. 2025</u>; <u>Nguyen, et al. 2022</u>; <u>Shi, et al. 2020</u>; <u>Yan, et al. 2014</u>). In this context, the significance of EBP lies not in serving as a simple substitute for traditional *Epimedium* medicinal materials but in representing a chemical system with its own compositional characteristics.

## 4. Conclusions

This study systematically characterized the chemical composition of *Epimedium sagittatum* bee pollen (EBP) by combining physicochemical analysis and UPLC–MS/MS-based targeted metabolomics, with Brassica rapa bee pollen (BBP), Camellia sinensis bee pollen (CBP), and *Epimedium* leaves used for comparison.

EBP showed a chemical profile clearly different from those of BBP and CBP. Its main feature was not the greatest metabolite diversity, but rather a relatively concen-trated flavonoid profile. EBP had the highest total flavonoid content among the three bee pollen types (4.75 mg/g), and flavonoid metabolites were the main contributors to its compositional separation from BBP and CBP. This pattern was reflected by several high-abundance unique metabolites, such as cacticin, brassicin, and tricetin-4′-methyl ether-3′-β-D-glucoside, as well as by 45 flavonoids that were consistently upregulated in EBP than RBP and CBP, including kaempferol, tamarixetin, and sinensetin. At the same time, although flavonoids were dominant in both EBP and *Epimedium* leaves, typical leaf constituents such as icariin and epimedins A–C were present at very low levels or were not detected in EBP. This difference suggests that bee pollen of tradi-tional medicinal plant chemical characteristics cannot be predicated only on the basis of the characteristic compounds of the source medicinal plant.

Taken together, EBP is better understood as a distinct food material with its own compositional features. Its most notable characteristic is a focused enrichment of fla-vonoid metabolites, unlike the leaf metabolite profile of Epimedium. Given the inte-gration of metabolomics and network pharmacology screening, flavonoid-related constituents, especially kaempferol, tamarixetin, and sinensetin, deserve particular at-tention in future studies of EBP. These findings provide a clearer compositional basis for understanding the chemical characteristics of bee pollen from traditional medicinal plants.

## Author Contributions

De-fang Niu: Conceptualization, Methodology, Writing - original draft, Data curation, Funding. Hai-xin Chen: Writing - original draft, Data curation. Cao-yang Lu: Methodology, Resources, Writing - review & editing. Cui-ping Zhang: Formal analysis, Writing - review & editing. Qiu-lan Zheng: Software, Super-vision, Investigation. Xiao-ling Su: Methodology, Writing - review & editing. Xiao-ming Fan: Formal analysis, Writing - review & editing. Yi-bo Luo: Formal analysis, Writing - review & editing. San-yao Li: Software, Supervision, Investigation. Bin Yuan: Formal analysis, Supervision, Writing - review & editing. Ping Liu: Software, Supervi-sion, Investigation, Funding. Fu-liang Hu: Supervision, Project administration, Fund-ing acquisition.

## Informed Consent Statement

Informed consent was obtained from all subjects in-volved in the study.

## Data Availability Statement

The data and analysis routines in this study are available on request.

## Supporting information

Supplementary Material for Review

