## Supplementary Material for Review for "Metabolomic characterization of *Epimedium sagittatum* bee pollen reveals a distinctive chemical profile": Supplementary document.docx

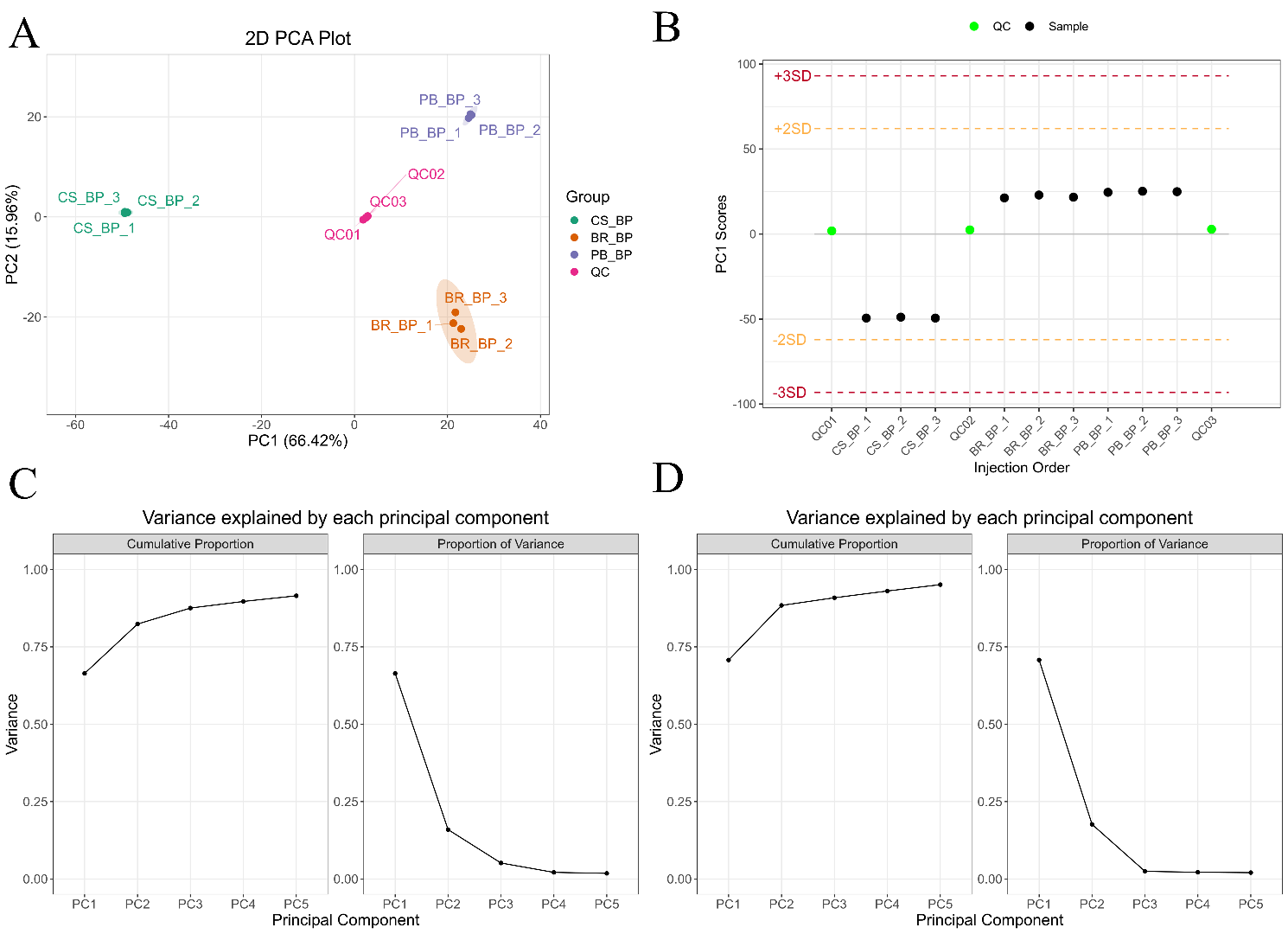

Figure S1(A)Overall sample PCA chart with QC. (B)Overall sample PC1 control chart. (C)Explanatory rate plot of the first 5 principal components of PCA for the overall sample. (D)Explanatory rate plot of the first 5 principal components of PCA for the overall sample with QC.

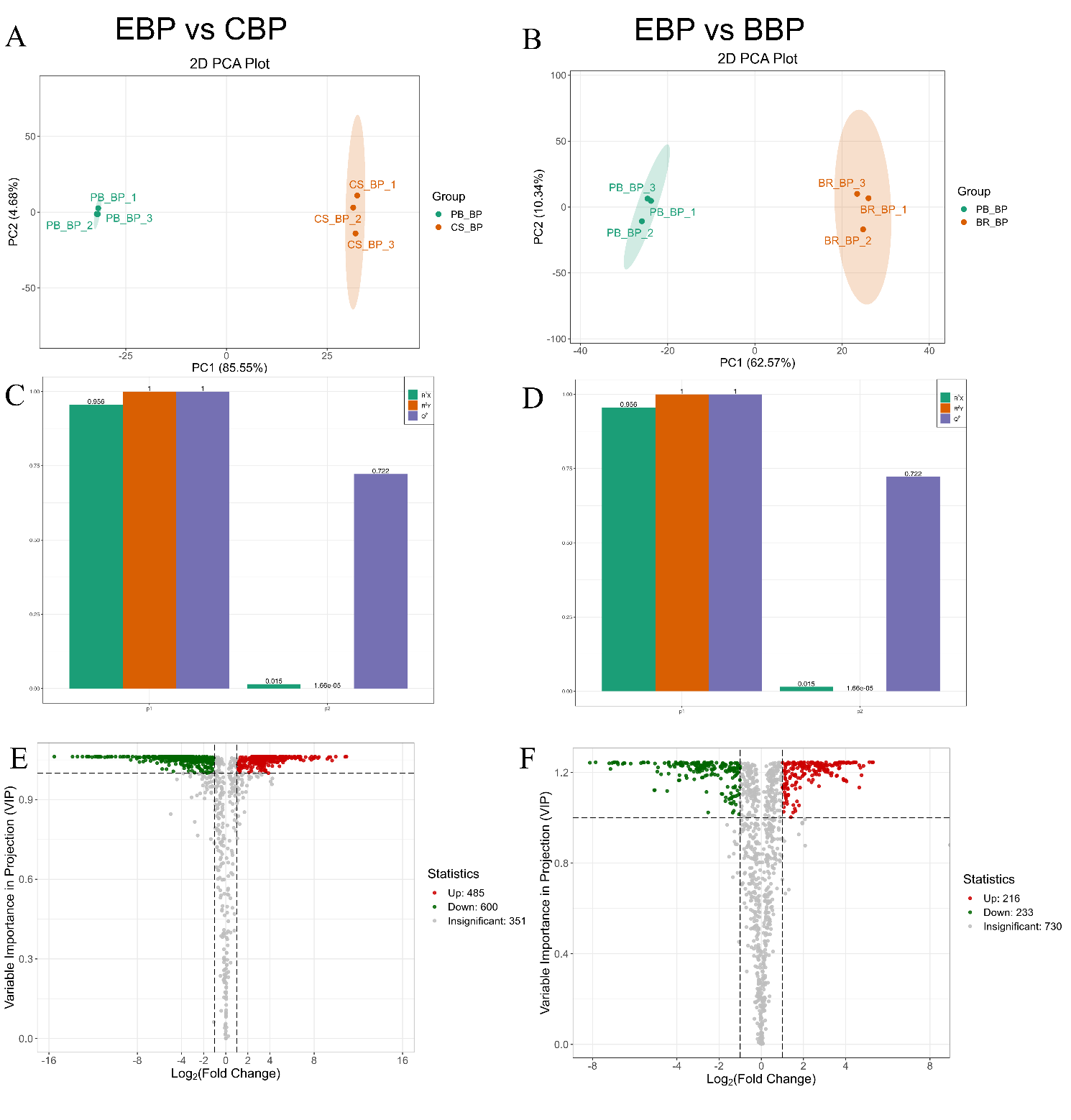

Figure S2 Metabolite PCA analysis plot of (A) EBP vs CBP group and (B)EBP vs BBP group. Overview diagram of the PLS-DA model for (C) EBP vs CBP and (D) EBP vs BBP. Volcano plot of differential metabolites in the (E) EBP vs CBP group and (F) EBP vs BBP group.

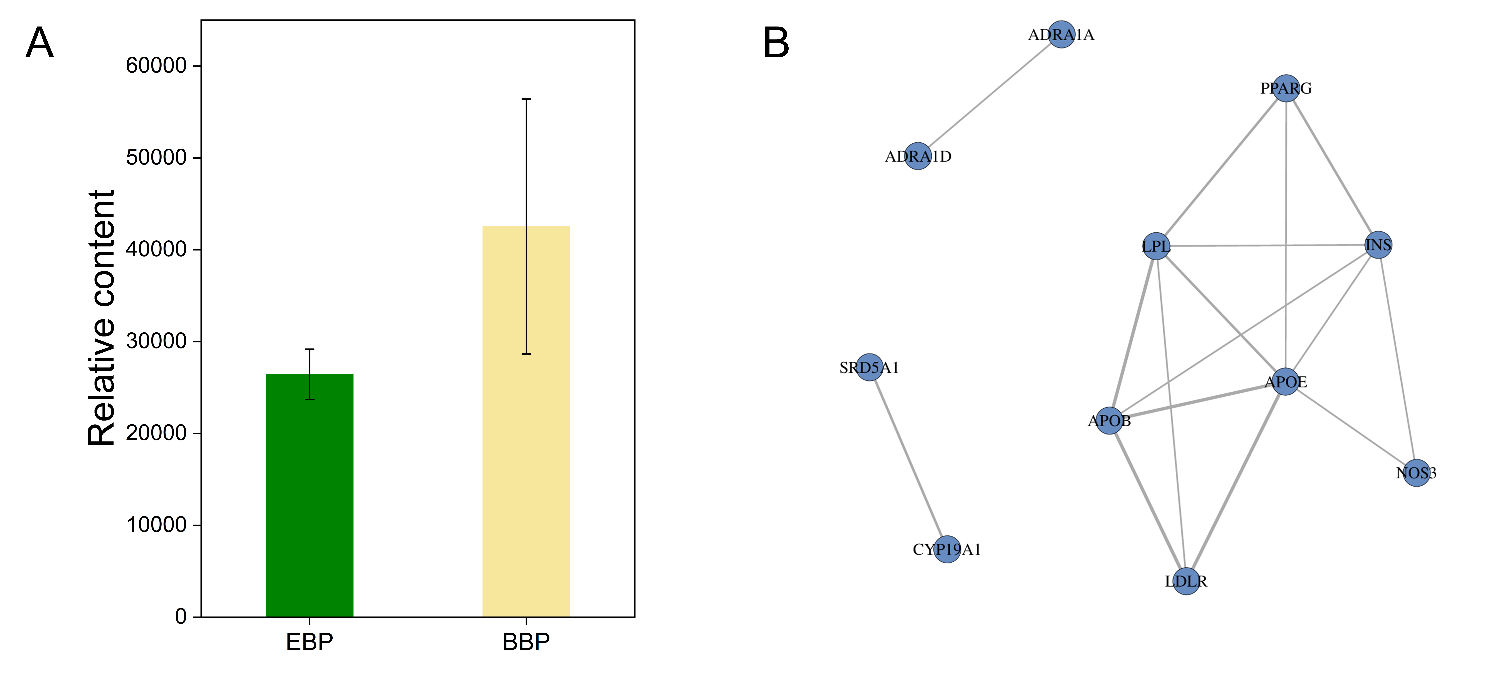

Figure S3 (A) Comparison of relative content of EBP vs BBP for *icariin.* (B) Target Interaction Network Diagram.
